# ProxiCapture Reveals Context-Dependent CRBN Interactome Landscape of Molecular Glue Degraders

**DOI:** 10.64898/2026.01.05.697692

**Authors:** Rubina Kazi, Henry J. Bailey, Jan Gerhartz, Varun Jayeshkumar Shah, Betül Toker, Julia Bein, Peter Wild, Radosław P Nowak, Thorsten Mosler, Ivan Dikic

## Abstract

Molecular glue degraders represent a rapidly expanding class of small molecules that reprogram E3 ubiquitin ligases to ubiquitinate and degrade disease-relevant proteins. Despite their therapeutic potential, the rational design of molecular glues remains challenging, underscoring the need for unbiased discovery strategies to identify new chemical targets. To address this challenge, we developed ProxiCapture, an affinity-based proteomics workflow that models the systemic behavior of molecular glues by combining purified CRBN-ΔHBD protein with native cell or tissue lysates. Systematic application of ProxiCapture across eight cancer cell lines, three maturation states of immune cells, and paired primary healthy and tumor tissues, revealed a comprehensive atlas of pomalidomide interactors, including previously uncharacterized targets. These findings reveal that degrader-dependent interactors of CRBN are context-dependent, requiring broad, physiologically and systemically anchored sampling to uncover the full “glueable” proteome. Taken together, this study establishes a scalable platform that accelerates molecular glue discovery by capturing cell- and tissue-specific recruitment profiles and predicting system-wide degrader effects.

## Introduction

Targeted protein degradation (TPD) has emerged as a transformative therapeutic strategy that harnesses the ubiquitin–proteasome system to eliminate disease-associated proteins from cells [1]. Two principal strategies dominate the TPD field: heterobifunctional degraders, or proteolysis-targeting chimeras (PROTACs)[2], and molecular glue degraders (MGDs)[3]. Both promote formation of a ternary complex between an E3 ubiquitin ligase and a target protein, inducing proximity-driven ubiquitination and subsequent proteasomal degradation [4,5].

MGDs are monovalent small molecules that reprogram E3 ligase specificity to recruit and degrade non-physiological substrates (“neo-substrates”), whereas PROTACs employ a bifunctional architecture that links a target-binding to an E3 ligase-recruiting ligand [6]. While PROTACs offer a modular and rational design, their large molecular size often leads to suboptimal pharmacokinetics, stimulating growing interest in molecular glues (MGs) as small-molecule-like modulators of E3 ligase specificity [7]. Unlike PROTACs, most of the MGs lack high-affinity binding to the target alone; instead, they stabilize weak or transient E3 ligase–substrate interactions, rendering rational discovery inherently difficult [8,9]. Indeed, the immunomodulatory drugs (IMiDs) thalidomide, lenalidomide, and pomalidomide were discovered serendipitously and only later identified as MGs that recruit neo-substrates to the CRL4-CRBN E3 ligase complex [10–12]. These MGDs target a structural glycine-containing beta-hairpin motif, commonly referred to as a “G-loop”, that fits into the thalidomide-binding pocket of CRBN and causes selective engagement of diverse substrates such as IKZF1/3, CK1α, and GSPT1[13]. Only recently, a non-β-hairpin motive was identified. The serine/threonine-kinase mTOR can be degraded, via a newly discovered glycine-containing helix-loop-helix (HLH) motive, adding non-β-hairpin proteins to the list of neo substrates[14]. All these compounds are today successfully used in clinical practice for the treatment of hematological malignancies such as multiple myeloma [15,16] and del(5q) myelodysplastic syndrome (MDS)[17,18]. Ongoing research aims to broaden the spectrum of cancers that can be targeted through MGs.

Global quantitative proteomics has been the standard approach to define molecular glue degradomes, using label-free or TMT-based quantification to track compound-dependent protein degradation [19–21]. Integration of Data Independent Acquisition (DIA) proteomics and ubiquitinomics has enabled large-scale screening, revealing that many MGDs act independently of the canonical G-loop degron [22]. However, degradome profiling remains incomplete, as these methods often miss transient, non-productive, or low-abundant ternary complexes, leaving much of the potential CRBN substrate space unexplored [8,23]. Proximity-labeling strategies such as BioID, TurboID, and AirID have addressed some of these gaps by fusing promiscuous biotin ligases to CRBN, enabling biotinylation of proteins that come within 10–30 nm during compound exposure [24–27]. While having the advantage of revealing protein-protein interactions *in-cellulo*, these approaches are limited by labeling efficiency, background noise, and scalability. Decades of medicinal chemistry around IMiD analogs led to the discovery of roughly 50 validated CRBN neo-substrates, most harboring a G-loop degron [28]. Yet, computational modeling across AlphaFold2 and Protein Data Bank structures predicts that CRBN-compatible G-loop geometries potentially exist in 1,633 proteins, including non-canonical binding configurations mediated by surface mimicry [14,29]. Thus, the complete CRBN substrate space remains poorly characterized.

IMiDs are FDA-approved MGDs that have mostly been characterized in cell–type–specific contexts. However, when administered to patients, they encounter diverse cellular and tissue proteomes, and their systemic molecular targets remain incompletely understood. We further hypothesized that the landscape of “glueable” substrates is highly context-dependent, shaped by cell identity, differentiation state, and the tissue microenvironment[22,30,31]. Since CRBN expression and substrate accessibility vary across biological systems, each degrader likely engages a unique subset of proteins defined by the biological system. Defining these context-dependent recruitment patterns is essential to anticipate the systemic effects, selectivity, and off-target liabilities of emerging MGs. To address this, we developed ProxiCapture, an affinity-based proteomics workflow that employs purified CRBN-ΔHBD incubated with native cell lysates to capture degrader-induced CRBN-protein interactions[32]. We employed pomalidomide, a clinically validated IMiD as a model MGD to interrogate the landscape of CRBN neo-substrates across three biological contexts: (1) cancer cell lines, (2) immune-cell differentiation states, and (3) normal and tumor tissues. This analysis yielded a comprehensive, system-wide pomalidomide interaction atlas encompassing 121 recruited proteins, including numerous previously uncharacterized and biological context-specific substrates. The atlas highlights how cellular composition and microenvironmental factors shape degradable substrate accessibility in complex systems.

## Results

### ProxiCapture reveals degrader-dependent CRBN interactors from native lysates

In our recent study [32], we developed an affinity-based proteomics strategy using a truncated recombinant human Cereblon (CRBN-ΔHBD) to explore structure–activity relationships among IMiD analogs. We showed that purified CRBN-ΔHBD retains full ligand-binding capacity and forms ternary complexes with canonical neo-substrates in cell lysates, confirming its suitability for degrader-interaction studies. Building on this foundation, we formalized a technological platform called ProxiCapture, which constitutes a robust proteomic workflow that allows systematic, comparative analysis of degrader-dependent CRBN-protein interactions across diverse biological contexts.

ProxiCapture is based on an immobilized N-terminally FLAG-tagged CRBN-ΔHBD that is exposed to native cell or tissue lysates in the presence or absence of an MG or PROTAC. Proteins that interact with CRBN in a degrader-dependent manner are enriched and analyzed by label-free quantitative mass spectrometry (Figure 1A). This robust workflow preserves endogenous protein complexes and enables the detection of both direct and secondary protein interactions with CRBN from nearly any biological matrix.

**Figure 1:**
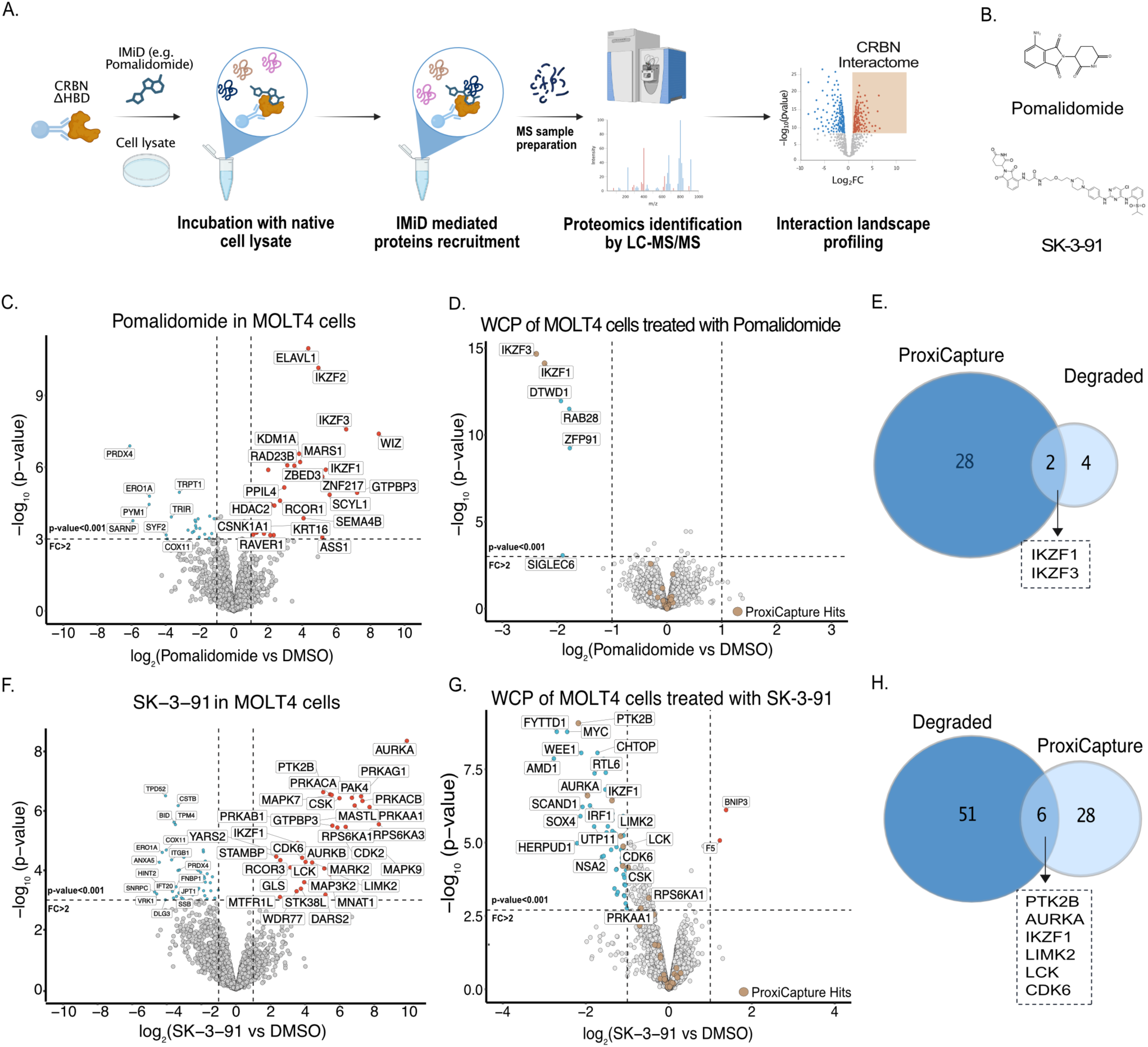
ProxiCapture: proof-of-concept using CRBN molecular glues and PROTACs. (A) Schematic of the ProxiCapture workflow. Purified N-terminally FLAG-tagged CRBN-ΔHBD was incubated with native lysate and CRBN-recruiting compounds, followed by anti-FLAG immunoprecipitation and label-free proteomic analysis. (B) Structures of the IMiD molecular glue pomalidomide and the CRBN-based multi-kinase PROTAC, SK-3-91. (C) Scatterplot depicting relative protein abundance following Flag-CRBN- ΔHBD enrichment (Data S1)from MOLT4 in-lysate treatment with 5μM pomalidomide with FC > 2, p < 0.001(left) and Scatterplot displaying relative protein abundance following treatment of MOLT4 cells for 5h with 1μM pomalidomide vs. DMSO with FC > 2, p < 0.001(right) (Data S1) (n = 3). (D) Venn diagram showing unique and overlapping protein hits comparing pomalidomide ProxiCapture and Global proteomics in MOLT4. (E) Scatterplot depicting relative protein abundance following Flag-CRBN- ΔHBD enrichment (Data S2)from MOLT4 in-lysate treatment with 5μM SK-3-91 with FC > 2, p < 0.001(left) and Scatterplot displaying relative protein abundance following treatment of MOLT4 cells for 5h with 1μM SK-3-91 vs. DMSO with FC > 2, p < 0.001(right) (Data S1) (n = 3). (F) Venn diagram showing unique and overlapping protein hits comparing SK-3-91 ProxiCapture and Global proteomics in MOLT4.

As a proof of concept, we applied ProxiCapture to MOLT4 cell lysates in the presence of the molecular glue pomalidomide (Figure 1B). The pomalidomide ProxiCapture profile (Figure 1C) revealed the canonical IMiD-based CRBN-protein interaction landscape, including IKZF1, IKZF3, WIZ, and CSNK1A1[11]. In addition to these known neo-substrates, ProxiCapture identified several proteins not previously reported as CRBN interactors, such as KDM1A, ELAVL1, and MARS1, as well as PPIL4 [29].

In addition to CRBN molecular glues, we wanted to test if ProxiCapture also resolves interactions that are induced by bifunctional molecules such as a PROTACs. We selected SK-3-91, a CRBN-based multi-kinase PROTAC, which was previously shown to degrade a broad section of kinases [21]. In contrast to pomalidomide, ProxiCapture in the presence of SK-3-91 PROTAC (Figure 1F) yielded a distinct set of proteins with a high enrichment for kinases, including AURKA, AURKB, CDK2, CDK6, MAP3K2, MARK2/3, PRKACA/B, and PAK4, consistent with the mode of action of SK-3-91 [19,21]. We observed an overlap between SK-3-91 and pomalidomide, which results from neo-substrate recruitment (IKZF1/3) induced by the CRBN-targeting component of SK-3-91. In line, we found co-enrichment of RNA-regulatory factors (ZBED3, ELAVL1, RCOR3, ZNF217) in the presence of SK-3-91. Together, these data showcase that ProxiCapture enables the reproducible detection of degrader-dependent CRBN-protein interactions across distinct chemotypes, supporting its use for systematic characterization of MGs and PROTACs.

To correlate recruitment to CRBN (ternary complex formation) with degradation outcomes, we performed global proteomic profiling of MOLT4 cells treated with pomalidomide or SK-3-91 (Figures 1D and 1G). We compared degrader-induced enrichment from ProxiCapture with protein degradation to distinguish non-productive from productive ternary complexes. Treatment with pomalidomide resulted in the selective depletion of canonical neo-substrates IKZF1, IKZF3, ZFP91, RAB28, and DTWD1, showing an overlap of 2 neo-substrates with ProxiCapture (Figures 1E). In contrast, the protein level of most newly captured interactors, including KDM1A and ELAVL1, was not altered, confirming their binding to CRBN without degradation. Conversely, SK-3-91 triggered broad degradation of kinase targets (AURKA, AURKB, CDK2 and others) along with IKZF1/3 via its CRBN-recruiting handle. Cross-comparison of the ProxiCapture and degradation datasets (Figure 1H) revealed an overlap of 6 kinases. In addition, we identified many CRBN-bound proteins that were not degraded, suggesting that ProxiCapture expands the CRBN target space, thereby complementing conventional degradation assays.

Together, these complementary enrichment and degradation datasets validate ProxiCapture as a sensitive approach for identifying degrader-induced CRBN recruitment events that remain invisible to global degradation profiling and supporting its use for systematic dissection of on-and off-target mechanisms of MGs and PROTACs.

### ProxiCapture reveals cell type-specific CRBN-protein interactions

We hypothesized that the spectrum of “glueable” substrates is unique to each cell type and highly dependent on the biological context. This diverse target space is determined by the individual proteome composition, differentiation status, and signaling networks characteristic of each biological system [22]. To assess this systemic diversity, we used the ProxiCapture to screen three biological contexts: (1) cancer cells, (2) distinct differentiation states of immune cells, and (3) tissues (Figure 2A).

**Figure 2.**
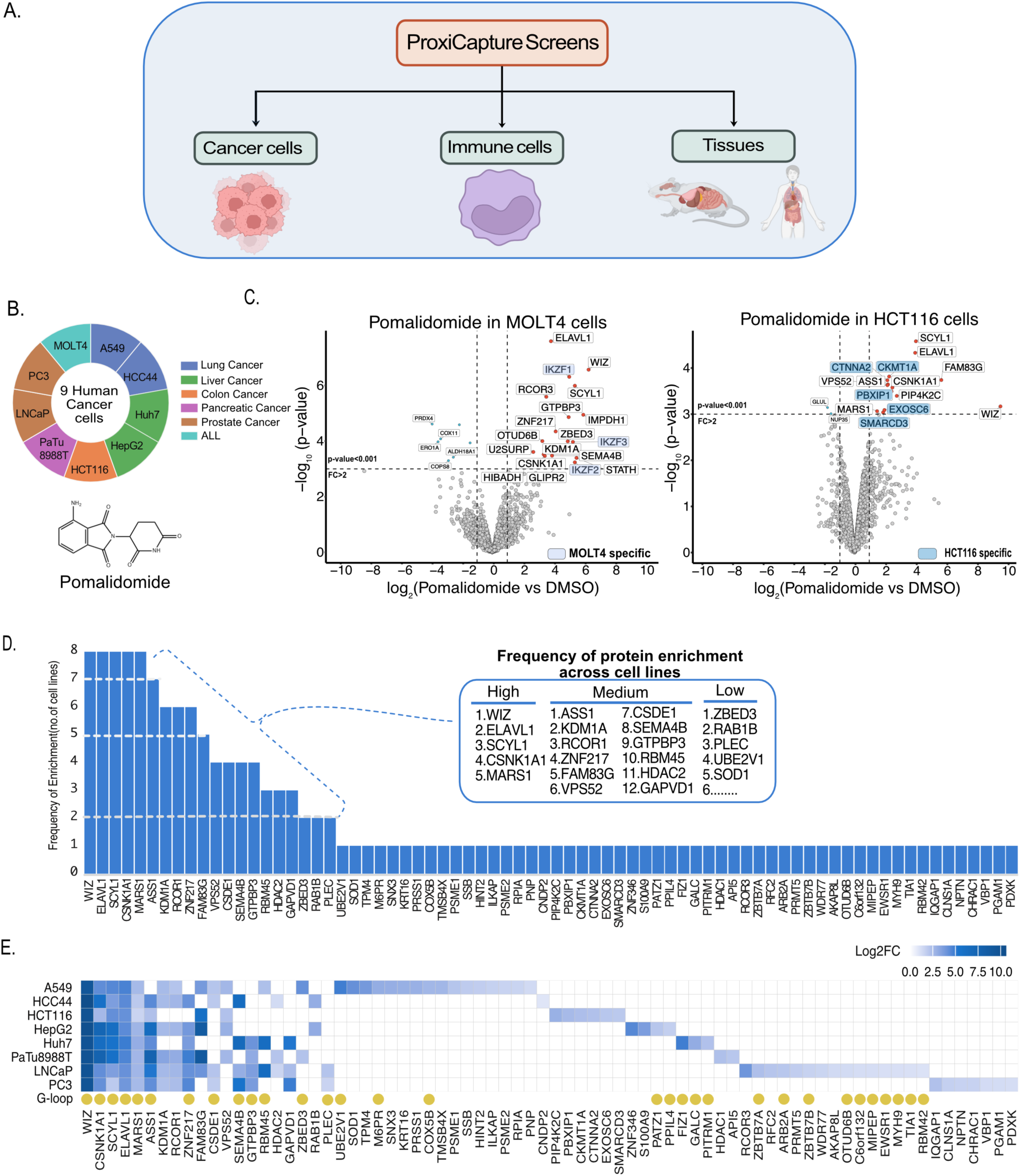
Cell-type-specific, pomalidomide-induced CRBN target space. (A) Overview of the three-tier screening design: cancer cell lines (n = 8), immune-cell states (n = 3), and tissues (mouse + human). (B) Donut chart showing six tumor types used for ProxiCapture. (C) Scatterplot depicting relative protein abundance following Flag-CRBN- ΔHBD enrichment from MOLT4 in-lysate treatment with 5μM Pomalidomide (left) and HCT116 in-lysate treatment with 5μM Pomalidomide (left)with FC > 2, p < 0.001(Data S2) (n = 3). (D) Frequency of enrichment across independent IP experiments for each identified target. The inset shows the top 20 proteins by enrichment frequency. (E) Heatmap showing log₂FC values of significant pomalidomide-responsive proteins (P < 0.001) in each cancer cell line. White space indicates log₂FC = 0 or no quantification. Statistical significance was assessed with a two-sided moderated t-test (FragPipe-Analyst).

Within the first context, we applied ProxiCapture to eight human cancer cell lines spanning five tumor types: lung (A549, HCC44), liver (HepG2, Huh7), colon (HCT116), pancreas (PaTu8988T), and prostate (LNCaP, PC3). We further included the MOLT4 leukemia model that is commonly used in IMiD and PROTAC discovery (Figure 2B, Supplementary Figure S1). These lineages collectively represent mechanistically diverse oncogenic programs and encompass tumor entities associated with high global incidence and mortality [33,34]. To assess the cell-dependent diversity of degrader-induced CRBN-protein interactions, we incubated each cell lysate individually with purified CRBN-ΔHBD in the presence of pomalidomide, enabling comparison of CRBN-protein interaction profiles across a large range of different tumor proteomes.

By applying ProxiCapture to MOLT4 cells, we were able to robustly identify IKZF1, IKZF2, and IKZF3, together with previously reported CRBN-interacting proteins ASS1, SCYL1, ZBED3, and ZNF217 [14,29]. This established a reference “hematopoietic” CRBN signature for benchmarking tissue-specific or lineage-divergent recruitment profiles. In contrast, colon-derived HCT116 cells displayed a distinct signature dominated by PBXIP1, a transcriptional co-activator upregulated in colon carcinoma [35], along with additional cell-specific interactors including CKMT1A, CTNNA2 [36], EXOSC6, and SMARCD3 [37] (Figure 2C). These results demonstrated that the same degrader can induce distinct CRBN-protein interactions across two different cell lines. This cell type-specific recruitment suggested to us that the substrate identification is highly influenced by the cellular proteome composition. By performing ProxiCapture across all mentioned tumor types, we were able to identify between 14 to 32 proteins per cell line that were recruited to CRBN in a pomalidomide-dependent manner (Supplementary Figure S1A). We further merged the hits from all datasets and found in total 74 CRBN-interacting proteins (Supplementary Figure S1B). Comparison between the MOLT4 reference and the hits from the cancer cell line screen revealed 11 shared proteins such as WIZ, KDM1A, and ELAVL1 and 63 unique proteins that engaged with CRBN (Supplementary Figure S1B), highlighting the need to screen a diverse range of cellular backgrounds for neo-substrate discovery.

Among the 74 combined hits from the cancer cell screen, we identified the known neo-substrates FAM83G, PATZ1, ZBTB7B, and MYH9 [28], whereas the well-established Ikaros zinc finger substrates were mostly enriched in our MOLT4 reference dataset (Supplementary Figure S1C). A frequency analysis across the eight cancer cell lines revealed only 20 proteins enriched in at least three backgrounds, including WIZ, ELAVL1, CSNK1A1, SCYL1, MARS1, KDM1A, RCOR1, ZNF217, FAM83G, and ZBED3 (Figure 2D). The remaining interactors appeared in only one or two cell types, reinforcing that pomalidomide-induced CRBN recruitment exhibits system-level variability driven by each unique cellular proteome landscape and signaling networks.

Mapping the log fold change of all enriched proteins across cell lines showed lineage-specific clustering, reflecting both tumor-intrinsic and system-wide differences in the CRBN substrate space (Figure 2E, Supplementary Figure S1D, E, F, G, H, I and J). Subsequently, we applied a recently developed method [29] in which the structural assessment software MASTER [38] is used to screen the AF2 Human genome library [39] for all proteins that contain a G-loop with a similar backbone (root-mean-squared-deviation (RMSD) cut off 1.5 Å) to the published G-loop degron of the well-known CRBN neo-substrate CK1llJ [18] (PDB: 5FAD, aa 35-42, 8 residues total, G at position 6). In addition, we screened for the recently described mTOR degron (PDB: 9NGT, aa 2087-2094, 8 residues total, G at position 6) [14]. The structural degron annotation identified 31 G-loop–containing proteins among the 74 total hits. While canonical β-hairpin motifs remain prevalent, nearly 60% of CRBN-recruited proteins lack a defined G-loop and likely engage through non-canonical or surface-mimicry interfaces (Figure 3E). This diversity suggests that CRBN recruitment extends beyond established degron rules, particularly in heterogeneous cancer contexts.

**Figure 3.**
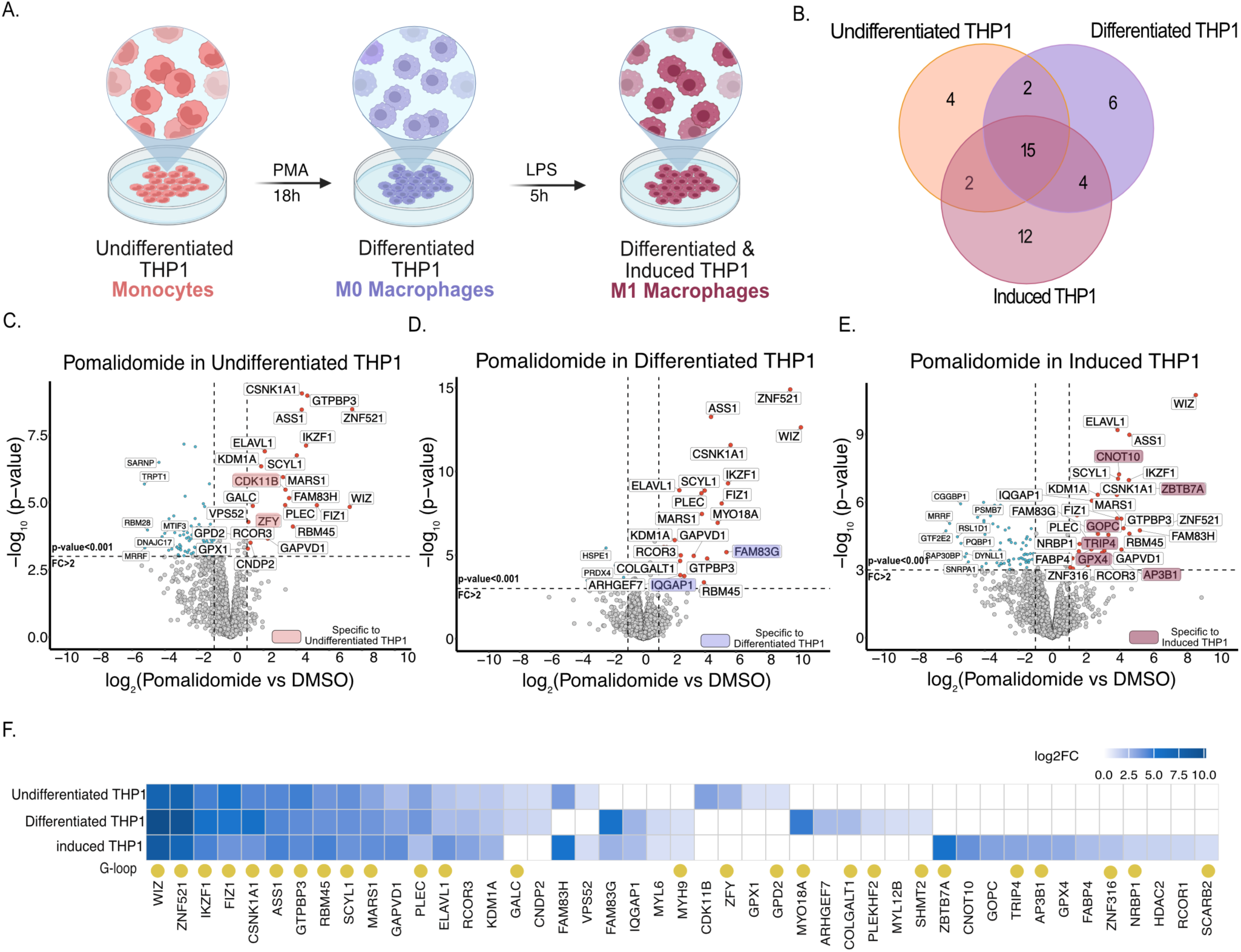
Immune state–specific CRBN interactome. (A) Differentiation scheme of THP-1 from monocytes into macrophages and structure of the pomalidomide used (B) Venn diagram showing unique and overlapping protein hits comparing ProxiCapture in undifferentiated, differentiated and induced THP1 cells. (C) Scatterplot depicting relative protein abundance following Flag-CRBN- ΔHBD enrichment from undifferentiated THP1 in-lysate treatment with 5 μM Pomalidomide with FC > 2, p < 0.001(Data S3) (n = 3). (D) As in C but for Differentiated THP1. (E) As in C but for Differentiated and induced THP1. (F) Heatmap showing log₂FC values of significant proteins (P < 0.001) in integrating three THP-1 states and five ligands showing clustering by immune-state identity. White space indicates log₂FC = 0 or no quantification. Statistical significance was assessed with a two-sided moderated t-test (FragPipe-Analyst).

### The CRBN interactome space depends on the cellular differentiation status

To examine how the cell differentiation state influences the degrader-induced CRBN protein interaction landscape, we next applied ProxiCapture to the THP-1 monocyte differentiation model. Immune differentiation and activation dramatically remodel proteomes and signaling pathways[40]. Thus, we tested whether CRBN associations change across immune maturation states. We generated three THP-1 states: undifferentiated monocytes, PMA-differentiated macrophages, and LPS-stimulated M1-like macrophages (Figure 3A) and applied ProxiCapture to lysates from each state using pomalidomide as the CRBN ligand. This setup models immune differentiation as a physiological axis of systemic variability and tests whether the same degrader engages distinct protein subsets along distinct differentiation trajectories.

The overall comparison between the THP-1 states (Figure 3B) revealed clear state-specific enrichment patterns, indicating that ProxiCapture sensitively detects dynamic CRBN recruitment during cellular differentiation. While we identified a core set of interactors that were common between all differentiation states (e.g., WIZ, ZNF521), we additionally found proteins that were unique to each stage (Figure 3C-E). In undifferentiated monocytes, CDK11B and ZFY were selectively enriched (Figure 3C), while the recruitment of IQGAP1 and FAM83G was favored in differentiated macrophages (Figure 3D). After LPS stimulation, which results in M1-like macrophages, CRBN was redirected towards transcriptional and trafficking regulators such as ZBTB7A, CNOT10, GOPC, TRIP4, and AP3B (Figure 3E). These unique patterns suggest that the CRBN interactor landscape is dynamically changing during immune maturation and highly depends on the cellular differentiation state.

To directly compare the THP-1 differentiation states, we assembled an enrichment matrix across the three conditions in the presence of pomalidomide (Figure 3F). Hierarchical clustering grouped samples by immune state, indicating that differentiation and activation status are the dominant determinants of CRBN recruitment when treated with the same molecule. Together, these findings show that even within a single immune lineage, pomalidomide-induced CRBN engagement varies systematically when the proteome is dynamically remodeled during immune polarization.

### Pomalidomide-induced CRBN-protein interactions are highly dependent on the tissue context

To assess whether ProxiCapture can resolve CRBN recruitment at the level of complex tissue proteomes, we next extended the workflow to primary mouse and human tissues (Figure 4A). CRBN is broadly conserved between human and mouse (∼96 % sequence identity; UniProt Q96SW2 vs Q91ZX7), and the thalidomide-binding pocket retains the same overall architecture. However, subtle residue substitutions within the thalidomide binding domain (TBD), notably near Val388 and His391, have been shown to alter neo-substrate recruitment efficiency [18,41,42]. Similar has been observed with the recruited substrate, for example mouse Sall4 contains key amino acid differences to human SALL4, which render it inactive for degradation by either mouse or human CRBN. Using human CRBN in mouse tissue lysates ensures that cross-species pulldowns reflect bona fide interactions rather than nonspecific binding. Mouse tissues were chosen to test the sensitivity of ProxiCapture in detecting subtle tissue-dependent differences under these homologous but non-identical conditions.

**Figure 4.**
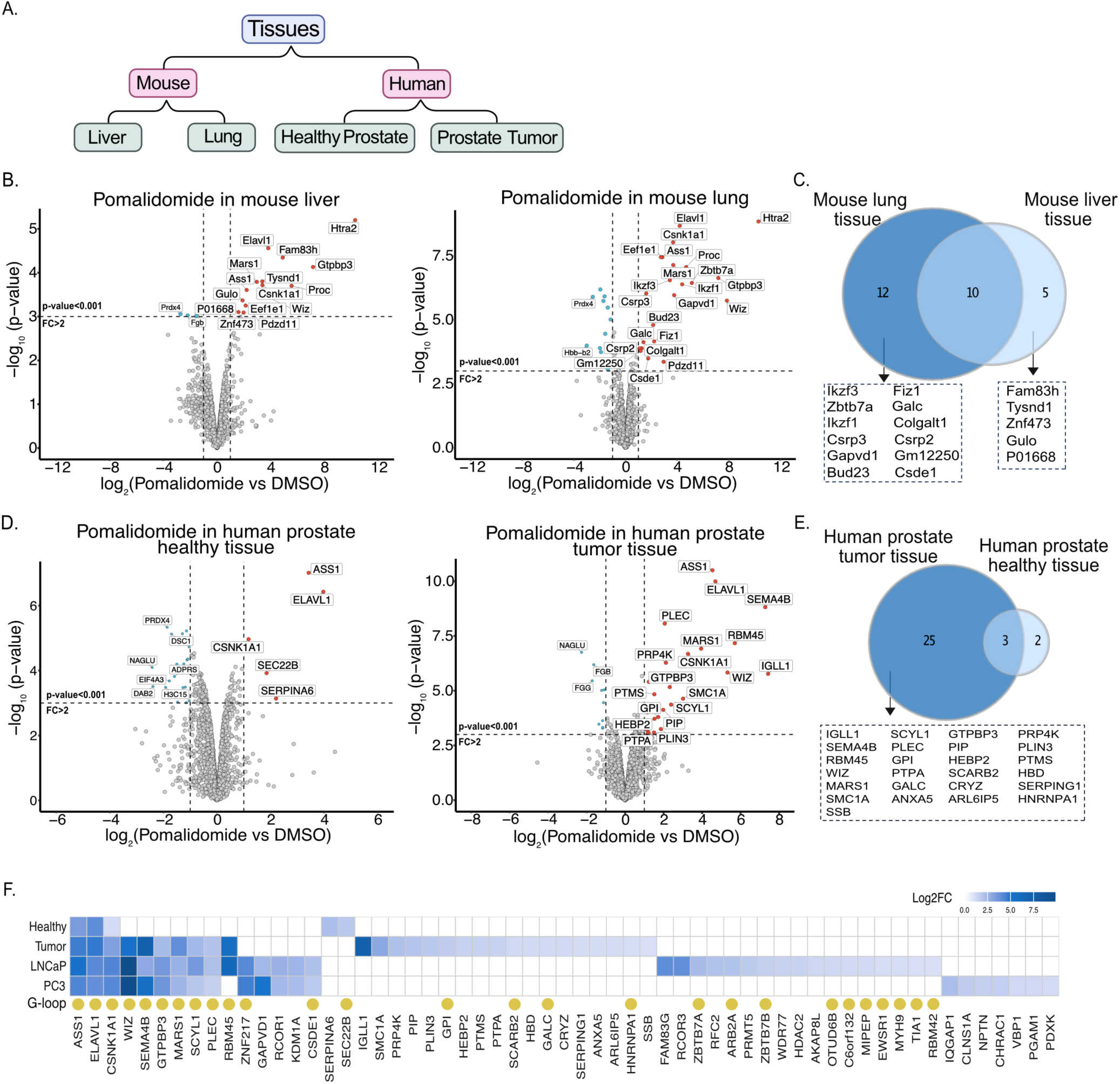
Tissue context–specific CRBN interactome. (A) Overview of tissue screen design. Human CRBN-ΔHBD and Pomalidomide were incubated with mouse (liver, lung) and human prostate (paired healthy/tumor) tissue lysates. (B) Scatterplot depicting relative protein abundance following Flag-CRBN-ΔHBD enrichment from mouse liver in-lysate treatment with 5μM Pomalidomide (left) and mouse lung in-lysate treatment with 5μM Pomalidomide (left)with FC > 2, p < 0.001(Data S4) (n = 3). (C) Venn diagram showing unique and overlapping protein hits comparing ProxiCapture in mouse liver lysate and mouse lung lysate. (D) Scatterplot depicting relative protein abundance following Flag-CRBN- ΔHBD enrichment from human healthy prostate tissue in-lysate treatment with 5μM Pomalidomide (left) and human tumor prostate tissue in-lysate treatment with 5μM Pomalidomide (left)with FC > 2, p < 0.001(Data S4) (n = 3). (E)Venn diagram showing unique and overlapping protein hits comparing ProxiCapture in human healthy prostate tissue and human tumor prostate tissue. (F) Heatmap showing log₂FC values of significant proteins (P < 0.001) in integrating two human prostate tissue and two human prostate cancer cells showing clustering by tissue and cell type. White space indicates log₂FC = 0 or no quantification. Statistical significance was assessed with a two-sided moderated t-test (FragPipe-Analyst).

ProxiCapture performed in murine liver and lung lysates in the presence of pomalidomide revealed distinct interaction profiles (Figure 4B). Notably, canonical CRBN substrates Ikzf1 and Ikzf3 were enriched exclusively in lung tissue, whereas neither was enriched from liver lysates, which instead contained metabolic and cytoskeletal proteins such as GalC, Csrp2, and Gm12250. Importantly, we didn’t observe Sall4 in these tissues either due to a lack of expression or the inability of human CRBN to recruit mouse SALL4[43]. The comparison between the two tissue types (Figure 4C) showed ten common neo-substrate proteins and a substantial fraction of unique proteins, showcasing that ProxiCapture can sensitively detect tissue-specific CRBN recruitment patterns from complex biological matrices.

Having established tissue-level resolution, we next applied ProxiCapture to human prostate tissues to examine CRBN engagement in a disease context. Matched healthy and tumor samples from the same donor were collected and analyzed (Figure 4D). Consistent with a more homogeneous cellular composition, the healthy prostate lysate yielded only a small set of pomalidomide-induced CRBN interactors. In contrast, the tumor tissue displayed markedly higher complexity, with 25 unique enriched proteins, including IGLL1, HBD, PLEC, PRP4K, and other secretory or immune-associated factors. These proteins are likely reflective of the remodeled tumor microenvironment and increased secretory activity characteristic of malignant prostate tissue (Figure 4E). Consistent with this observation, whole-cell proteomics of the same matched healthy and tumor prostate tissues (Supplementary Figure S2A) revealed extensive proteome remodeling, with numerous proteins significantly upregulated or downregulated in the tumor sample, including inflammatory, secretory, and cytoskeletal factors. Importantly, ProxiCapture enrichment does not simply mirror protein abundance, since the CRBN interactors identified in tumor tissue include proteins that are upregulated, downregulated, or unchanged at the whole-proteome level.

To relate tissue-derived interactomes to cellular models, we compared prostate tumor–tissue interactors with those identified in prostate cancer cell lines (LNCaP and PC3) (Supplementary Figure S2B). The 18 shared hits define a core set of pomalidomide-induced CRBN-protein interactions found in prostate cancer cells, whereas tissue-specific interactors likely originate from stromal or microenvironmental compartments that are absent in cultured cell lines. A comparative heatmap integrating tissue and cell datasets (Figure 4F) clustered samples by biological source rather than by individual protein identity, suggesting that CRBN accessibility is shaped primarily by tissue context and multicellular architecture. Degron annotation classified captured proteins into canonical G-loop and non-G-loop groups, confirming that both structural classes contribute to the tissue-resolved CRBN interactome and to systemic heterogeneity in degrader response. Because some G-loop–dependent degrons are not fully conserved between mouse and human, these interactions may show species-specific behavior; ProxiCapture therefore provides a practical way to verify which animal models are compatible with a given molecular glue or PROTAC target before advancing to in vivo studies.

### Comprehensive analysis of the three-tier ProxiCapture screens

For further downstream analyses, we integrated all datasets obtained from the three individual biological tiers to classify CRBN interactors by degron architecture, distinguishing canonical β-hairpin G-loop and non-G-loop proteins through structural motif annotation. We have followed a method recently developed by Baek *et al* [29] where structural assessment software MASTER [38] was used to screen the AF2 Human genome library[39] for all protein containing a g-loop with a similar backbone (root-mean-squared-deviation (RMSD) cut off 1.5 Å) to the published g-loop degron of the well-known CRBN neo-substrate CK1llJ [18] (PDB: 5FAD, aa 35-42, 8 residues total, G at position 6). The same was repeated for the mTOR degron, recently described (PDB: 9NGT, aa 2087-2094, 8 residues total, G at position 6) [14]. In total, this analysis identified 49 proteins containing a G-loop–like motif and 83 proteins lacking this architecture, providing a clear classification of CRBN interactors based on degron structural features.

To obtain a global view of CRBN recruitment and evaluate how degrader responses integrate across biological systems, we merged all significant interactors (log fold change > 1, p < 0.001) from the cancer-cell, immune-cell, and tissue ProxiCapture screens, generating a non-redundant set of 121 proteins. Of those proteins, 20 turned out to be very frequent hits independent of the biological background, with the most frequent being ELAVL1, CSNK1A1, and WIZ (Figure 5B). In addition to the well-established CRBN substrates and interactors carrying β-hairpin G-loop degrons, we identified many proteins that have not yet been shown to interact with CRBN. Importantly, more than half of the proteins appeared only in one or two screens, underscoring the system-level variability of CRBN recruitment and highlighting the importance of charting a diverse biological space when screening for novel molecular glue targets.

**Figure 5.**
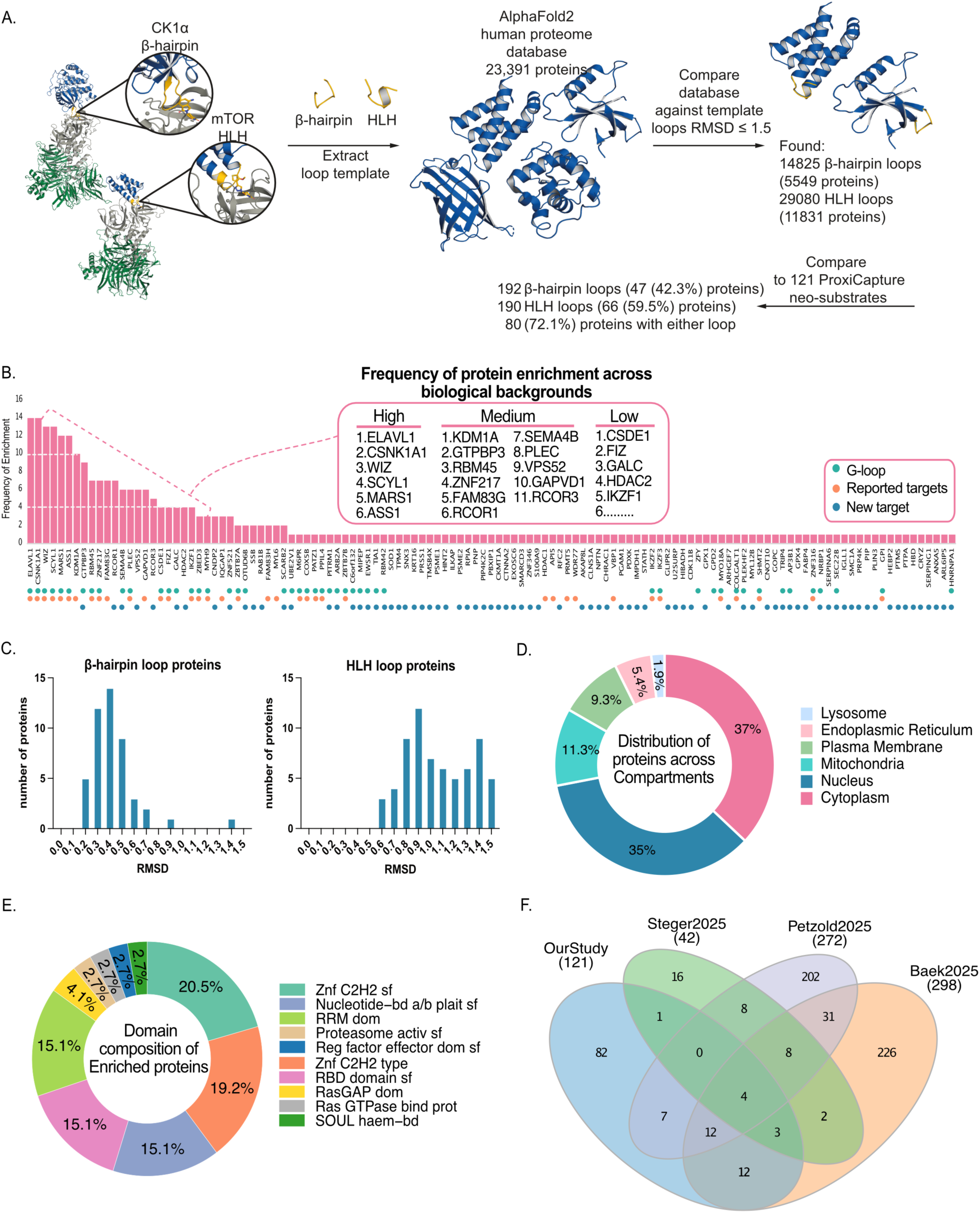
Integrated analysis across cells, immune states, and tissues. (A) MASTER analysis workflow (B) Frequency distribution of significant CRBN-interacting proteins (log₂ FC > 1, P < 0.001) identified across cancer-cell, immune-cell, and tissue ProxiCapture screens. Known CRBN neosubstrates, proteins containing β-hairpin or helical G-loop degrons, and newly identified targets are highlighted (Data S5). (C) Structural loop-motif classification of the 121 ProxiCapture-enriched proteins based on degron features (Data S6). (D) Donut plot showing subcellular localization distribution based on GO:CC terms retrieved via *biomaRt* (Ensembl v111) (E) Donut plot showing domain-superfamily composition from InterPro database using DAVID. (F)Venn Diagram comparing ProxiCapture interactome with recent CRBN-target studies (*Petzold et al.*, 2025; *Baek et al.*, 2025; *Steger et al.*, 2025) showing shared canonical substrates and novel context-specific interactors unique to this study (Data S5).

To explore the structural features underlying these interactions, we performed loop-motif analysis of all 121 proteins using degron annotation criteria as previously described[14,29] (Figure 5C, Supplementary Figure S3A, B, C). All proteins containing a CK1llJ-like β-hairpin loop with a Gly at position 6, were classified as G-loop-containing, whereas proteins with a similar backbone but lacking Gly at position 6, or those without any comparable loop-motive were considered non-G-loop containing. Proteins containing only an HLH loop were also placed into the non-G-loop category. Our analysis revealed that 42.3% of all 121 proteins contain a CK1llJ-like β-hairpin loop. Examination of the RMSD distribution (Figure 5C) revealed that most β-hairpin-like loop proteins already met the filter criteria for low RMSD (<0.8). Higher RMSD (>0.8) did not substantially increase the total amount of G-loop containing proteins. This stands in contrast to the HLH mTOR-like degron, for which the first hits did not only exhibit higher RMSDs to the mTOR degron, but also the number of prospective hits increased with the RMSD cut-off. This observation emphasises the robustness of the β-hairpin loop degron, promoting its application for future CRBN neo-substrate identification.

Next, we examined the subcellular distribution of the 121 proteins using Gene Ontology Cellular Component (GO:CC) terms retrieved via *biomaRt* (Ensembl v111) (Figure 5D). Approximately 37 % of interactors localize to the cytoplasm and 35 % to the nucleus, with smaller fractions associated with mitochondria (11 %), endoplasmic reticulum (9 %), and plasma membrane (5 %). This pattern likely reflects the accessibility of CRBN-bound complexes within soluble fractions. We note that lysate-based assays can also reveal interactions between proteins that would normally not meet in the cell, although the broad expression of CRBN helps to limit this concern. Overall, the data remain consistent with the reported localization of CRL4^CRBN^ and IMiD-responsive substrates[22,44].

To evaluate domain composition, we retrieved annotations from the InterPro database in DAVID[45,46] and grouped the proteins at the superfamily and family levels (Figure 5E). The most represented class was the C2H2 zinc-finger superfamily (20.5 %), followed by C2H2 zinc-finger type (19.2 %), nucleotide-binding α/β plait superfamily (15.1 %), RNA-recognition motif (RRM) (15.1 %), and Ras GTPase-binding domain proteins (15.1 %). These distributions are parallel to previously reported IMiD-dependent target profiles dominated by zinc-finger transcription factors and RNA-binding proteins [14,29]. The recurrent enrichment of nucleic-acid-binding domains across cancer, immune, and tissue datasets points to shared systemic nodes of CRBN accessibility beyond classical degron classes.

To explore biological processes, we performed Gene Ontology enrichment using clusterProfiler [47] (Supplementary Figure S3D). Top 15 Significantly enriched terms (p.adjust < 0.05) included mononuclear-cell differentiation, positive regulation of cell development, mRNA processing, pyridine-containing compound metabolic process, autophagosome assembly, and stress-granule assembly. Together, these functions indicate that CRBN neosubstrates recruited by molecular glues relate to transcriptional control, RNA metabolism, and stress-response circuitry across contexts.

We generated a STRING functional association network (Supplementary figure S3E), which resulted in several densely connected clusters corresponding to chromatin regulators (KDM1A, RCOR1, HDAC1), zinc-finger transcription factors (IKZF1/3, ZNF217, ZBTB7A), RNA-binding proteins (ELAVL1, GTPBP3, RRM-containing factors), and signaling molecules (CSNK1A1, SCYL1). These pathway-linked groupings, combined with numerous singleton nodes, support that ProxiCapture recovers true CRBN-proximal substrates, and that enrichment is driven by ligand-dependent recruitment.

Finally, we compared our data obtained by ProxiCapture with three recent studies that charted CRBN-protein interactions by different approaches [14,22,29] (Figure 5E). We found that 39 proteins were shared across all datasets, validating assay specificity and suggesting a core set of proteins that is recruited to CRBN in an IMiD-dependent manner. Our study revealed an additional 82 unique CRBN interactors, underlining the importance of screening a diverse range of biological systems. Thus, beyond expanding the known CRBN target space, our recruitment-centric approach recovers non-degradative interactors alongside degradable substrates, enabling system-level prediction of on- and off-target engagement. Moreover, we identified cancer type-specific CRBN targets, suggesting that pomalidomide and related derivatives may have therapeutic potential in diseases beyond classical hematological malignancies.

## Discussion

The discovery of molecular glues has transformed targeted protein degradation by showing how small molecules can remodel E3 ligase-substrate interfaces to achieve selective proteome manipulation [28]. Yet discovery remains largely empirical because of the lack of systematic workflows that capture context-dependent and low-abundant interactions [7]. ProxiCapture addresses this gap by directly detecting degrader-induced CRBN recruitment from native lysates and thereby expanding the known CRBN target space across cells and tissues. Rather than inferring engagement from steady-state protein loss, ProxiCapture reads out binding events themselves, thereby encompassing both degradative substrates and non-degradative (“bystander”) interactions that may still influence complex stability, localization, or signaling topology. Because it operates in native proteomes without genetic manipulation, the approach preserves endogenous stoichiometry and is readily transferable across diverse biological matrices, enabling systematic, cross-context mapping of CRBN recruitment.

Our data show that the CRBN recruitment landscape is strongly shaped by biological context. Across eight cancer cell lines spanning five tumor types, the pomalidomide-induced interactome exhibited lineage-specific signatures with limited overlap, consistent with cell type–specific expression, differential post-translational modification, and distinct accessibility of degron-containing proteins. In the immune system, differentiation of THP-1 monocytes into macrophages and acquisition of an M1-like state reconfigured CRBN recruitment, revealing activation-dependent changes in substrate availability. Tissue analyses extended this principle: mouse liver and lung lysates yielded distinct interactomes despite identical CRBN sequences [18,42], and human prostate tumor tissue displayed broader engagement than matched normal, underscoring that proteomic context, rather than E3 sequence, dominates recruitment. Together, these observations argue against a universal “CRBN degradome” and instead support a model in which substrate accessibility is dynamically governed by proteomic composition, cellular state, and signaling architecture.

Integration of all datasets defined 121 high-confidence interactors, enriched in chromatin regulators, RNA-binding proteins, and signaling enzymes. Many of these are non-canonical with respect to established G-loop substrates, indicating that degron geometry alone does not dictate recruitment; rather, contextual accessibility and complex assembly shape the glueable proteome. The presence of numerous non-degradative interactors suggests that MGs can exert functions beyond targeted depletion, transiently remodeling protein networks and potentially influencing RNA metabolism, stress responses, or differentiation programs. In that vein, newly observed factors bridging chromatin regulation, RNA processing, and stress signaling, including FAM83G, ZNF217 and ELAVL1, illustrate how CRBN engagement can intersect with broader regulatory nodes that may not manifest as immediate degradation but could still modulate cellular behavior.

Recent studies have shed first light on the IMiD-induced CRBN-protein interaction landscape[14,22,29]. While using distinct methods to look at CRBN interactions (proximity labeling, co-IP, in silico prediction), the overlap of the studies suggests a core set of proteins that is recruited to CRBN independent of the technical approach. However, previous studies have rather focused on diversifying the IMiD chemistry rather than exploring a larger biological space. A central outcome of this study is the expansion of the CRBN target space, with 82 previously uncharacterized interactors uncovered by ProxiCapture. Although our experiments focus on a single IMiD scaffold (pomalidomide), applying the same workflow to a broad range of biological contexts shifted the limit closer to the theoretical upper bound of CRBN-recruitable substrates. Moreover, we identified many novel cancer type-specific interactors of CRBN, suggesting that pomalidomide and close derivatives bear therapeutic potential in other malignancies beyond hematological cancers. The chemical generalizability, coupled with the ability to run in native proteomes from primary cells or tissues, positions ProxiCapture as a powerful entry point for degron discovery and for prioritizing targets and chemotypes tailored to specific disease contexts.

### Implications and future directions

ProxiCapture reveals that CRBN recruitment is highly dependent on the biological context, arguing against a universal “CRBN degradome” and underscoring the need to map degrader effects across multiple cell types and tissues to anticipate efficacy and safety. Next steps include adding temporal perturbations (e.g., hypoxia, stress, differentiation) and patient-derived samples to define how disease state reshapes “glueable” proteomes. The framework is readily extendable beyond CRBN to other E3 ligase complexes (VHL, DCAF15, RNF114 and others with reported small molecule binders) and to proximity-inducing chemistries without degradation. Integrating ProxiCapture with quantitative ubiquitinomics, crosslinking MS, and machine-learning-based–based degron prediction will enable proteome-scale models linking recruitment to outcome In sum, ProxiCapture transforms molecular glue discovery into a systematic and system-aware exploration of ligase–substrate space. By bridging biochemical precision with physiological and tissue-level relevance, this platform provides a foundation for predicting the systemic effects of molecular glues and for uncovering new therapeutic nodes within the human degradable proteome.

### Limitations of the study

Although ProxiCapture adds substantial new resolution to CRBN recruitment events, the assay alone cannot provide a complete picture of degrader function, particularly with respect to downstream ubiquitination, ternary-complex dynamics, or cellular degradation outcomes. Accordingly, ProxiCapture is most powerful when integrated with complementary assays within a broader screening platform to build a holistic view of degrader behavior.

The computational MASTER analysis only relies on structural similarity of the g-loop while not taking surrounding structures into consideration. It is to be expected that a clash score analysis of the identified g-loops in the CRBN binding pocket would reduce the number of g-loop containing proteins.

## Materials and Methods

### Materials

Pomalidomide and SK-3-91 were purchased from Merck Healthcare KGaA. A549, HCC44, Huh7, HepG2, HCT116, and PaTu8988T cells were maintained in DMEM medium (Gibco, Life Technologies) supplemented with 10% fetal bovine serum (FBS) (Gibco, Life Technologies) and 1% penicillin/streptomycin (Gibco, Life Technologies). LNCaP cells were maintained in EMEM medium (Cytion) supplemented with 10% fetal bovine serum (FBS) (Gibco, Life Technologies) and 1% penicillin/streptomycin (Gibco, Life Technologies). PC3 cells were maintained in DMEM/F-12 medium (Gibco, Life Technologies) supplemented with 10% fetal bovine serum (FBS) (Gibco, Life Technologies) and 1% penicillin/streptomycin (Gibco, Life Technologies). MOLT4 cells were a kind gift from the Winter lab, AITHYRA, AT and were maintained in RPMI 1640 (Gibco, Life Technologies) supplemented with 10% fetal bovine serum (FBS) (Gibco, Life Technologies) and 1% penicillin/streptomycin (Gibco, Life Technologies).

### THP1 Differentiation

THP-1 wild-type cells (ATCC) were prepared under three conditions: (i) undifferentiated; (ii) PMA-differentiated; and (iii) PMA-differentiated followed by LPS priming. For differentiation, cells were treated with 100 ng/mL phorbol 12-myristate 13-acetate (PMA; MedChem Express, HY-18739) for 18 h at 37 °C, 5% CO . Where indicated, PMA-differentiated cells were subsequently primed with 0.15 µg/mL lipopolysaccharide (LPS; InvivoGen, O111:B4) for 5 h.

After treatment, cells were washed once with Dulbecco’s phosphate-buffered saline (DPBS; Gibco, 14190144) and detached with 0.05% trypsin-EDTA (Gibco, 25300054). Residual trypsin was removed by centrifugation (700 × g, 5 min, 4 °C) followed by two DPBS washes. Finally, cells were lysed in IP buffer (50 mM Tris-HCl, pH 7.5; 120 mM NaCl; 1% NP-40; 0.5 mM EDTA; protease inhibitors; phosphatase inhibitors; and N-ethylmaleimide [NEM]). Lysates were sonicated in a bath sonicator for 5 cycles (10s on / 30s off) at 4 °C. After clarification by centrifugation, supernatants were transferred to low-bind tubes and used for ProxiCapture.

### Tissue lysate for Flag-CRBN IP

Tissue samples were obtained as follows: five healthy and five tumor prostate tissues from matched donors, and liver and lung tissue from a 3-month-old wild-type mouse. Samples were homogenized with disposable pestles and lysed in IP buffer (50 mM Tris-HCl, pH 7.5; 120 mM NaCl; 1% NP-40; 0.5 mM EDTA; protease inhibitors; phosphatase inhibitors; and N-ethylmaleimide [NEM]). Lysates were sonicated in a bath sonicator for 10 cycles (10s on / 30s off) at 4 °C. After clarification by centrifugation, supernatants were transferred to low-bind tubes and used immediately for ProxiCapture. Lysate from five human prostate tissues were pooled together and performed ProxiCapture in technical triplicates.

### ProxiCapture: Flag-CRBN IP

Purified Flag-CRBN was conjugated to Flag beads (cat no. A36797, Thermofischer) in IP buffer (50mM Tris pH-7.5, 120mM NaCl, 1% NP40, 0.5mM EDTA) for 1hr at 4^0^C on a rotating shaker, before addition of DMSO or 5µM Pomalidomide/SK-3-91 for 30min at 4^0^C. Subsequently, ∼1mg of freshly prepared cells of tissue protein lysate was added to the prepared beads in IP lysate buffer (50mM Tris pH-7.5, 120mM NaCl, 1% NP40, 0.5mM EDTA, protease inhibitors, phosphatase inhibitors and NEM) for 2hr at 4^0^C while rotating. The beads were washed three times with IP buffer and afterwards used either for western blotting or trypsin digestion followed by LC-MS^2^ analysis.

### Mass spectrometry sample preparation (ProxiCapture)

Protein-bound flag-CRBN beads were incubated with 20 µl SDC buffer (3% sodium deoxycholate in 50 mM Tris-HCl pH 8.5) and heated for 5min at 65°C and supernatant was collected. This step was repeated one more time to get elute of 40µl. Further, reduction and alkylation were performed using 5ul of 5mM TCEP, 20mM CAA in 50 mM Tris-HCl pH 8.5 at 95°C for 10min. 500 ng of trypsin in 45 µl 50 mM Tris-HCl (pH 8.5) was added to each sample and kept it for digestion at 37 °C overnight. The digestion was stopped upon addition of 135 µl of 1% TFA in isopropanol. Peptide clean-up was performed using SDB-RPS stage tips (Sigma-Aldrich). Peptides were added to stage tips and washed first with 1% TFA in isopropanol and then with 0.2% TFA in water. Lastly, peptides were eluted in 80% acetonitrile plus 1.25% ammonia and dried in vacuum concentrator.

### Mass spectrometry data acquisition (ProxiCapture)

Dried peptides were resuspended in 2%ACN with 0.1% TFA and used for LC-MS^2^ analysis on a QExactive HF mass spectrometer coupled to an easy nLC 1200 (Thermo Fisher Scientific) fitted with a 35 cm long, 75µm ID fused-silica column packed in house with 1.9 µm C18 particles (Reprosil pur, Dr. Maisch). The column was maintained at 40 °C using an integrated column oven (Sonation). Peptides were eluted in a non-linear gradient of 5–40% acetonitrile over 60 min and sprayed into the mass spectrometer equipped with a nanoFlex ion source (Thermo Fisher Scientific). Full-scan MS spectra (300–1,650 *m*/*z*) were acquired in profile mode at a resolution of 60,000 at *m*/*z* 200, a maximum injection time of 20 ms and an AGC (automatic gain control) target value of 3 × 10^6^. Up to 10 of the most intense peptides per full scan were isolated using a 1.4-Th window for fragmentation by higher energy collisional dissociation (normalized collision energy of 27). MS/MS spectra were acquired in centroid mode with a resolution of 30,000, a maximum injection time of 54 ms and an AGC target value of 1 × 10^5^. Single charged ions, ions with a charge state of more than seven and ions with unassigned charge states were not considered for fragmentation, and dynamic exclusion was set to 20 s to minimize the acquisition of fragment spectra representing already acquired precursors.

### Mass spectrometry data analysis (ProxiCapture)

MS raw data were analyzed using FragPipe v21.1, with MSFragger v.4.0 [48] and Philosopher v.5.1.0 [49]. The built-in workflow “LFQ-MBR” was used with a precursor mass tolerance of 20 ppm and fragment mass tolerance of 20 ppm. The human proteome database used by FP (ID: UP000005640, 02/02/2025) comprised of 20,454 reviewed sequences only and their corresponding decoys, including common contaminant proteins. Identifications were filtered to obtain false discovery rates (FDR) below 1% for both peptide spectrum matches (minimum peptide length of 7) and proteins using a target-decoy strategy. For all searches, carbamidomethylated cysteine was set as a fixed modification and oxidation of methionine and N-terminal protein acetylation as variable modifications with allowing up to 3 modifications per peptide. Strict trypsin cleavage was set as protein digestion rule. Label-free quantification was performed using IonQuant v.1.10.27[50]. Data were further processed using FragPipe Analyst [51]. Subsequently, the data were plotted in R using custom scripts.

### Mass spectrometry sample preparation proteome analysis

Cells were treated with DMSO and 1μM Pomalidomide/ SK-3-91 in biological triplicates for 5hr. Cells were harvested and washed with PBS by centrifugation. Further, Lysis buffer (2% SDS, 50mM Tris pH 8.5, 10mM TCEP, 40mM CAA, supplemented with protease inhibitor cocktail) was added to the pellets and cells were homogenized by sonication in ice and boiling at 95 °C for 10 min. Proteins were precipitated using methanol-chloroform and resuspended in 8 M urea, 50 mM Tris pH 8.5. Bradford assay was used to determine final protein concentration in the lysate. 50ug of proteins were digested with 1:50 w/w LysC (Wako Chemicals, cleaves at the carboxylic side of lysine residue) and 1:100 w/w trypsin (Promega, Sequencing-grade) overnight at 37 °C after dilution to a final urea concentration of 1 M using 50 mM Tris pH 8.5. Digested peptides were then acidified (pH 2–3) using trifluoroacetic acid (TFA) and purified using C18 SepPak columns (Waters). Desalted peptides were dried and resuspended in TMT-labeling buffer (200 mM EPPS pH 8.2, 20% acetonitrile). 10μg of peptides per condition were subjected to TMT labeling with 1:2.5 peptide TMT ratio (w/w) for 1 h at room temperature. The labeling reaction was quenched by addition of hydroxylamine to a final concentration of 0.5% and incubation at room temperature for 15 min. Successful TMT labeling was verified by mixing equimolar ratios of peptides and subjecting the mix to single shot LC-MS/MS analysis. For high pH reversed phase fractionation on a Dionex analytical HPLC, 50µg of pooled and purified TMT labelled samples were resuspended in 10mM ammonium-bicarbonate (ABC), 5%ACN, and separated on a 250mm long C18 column (Aeris Peptide XB-C18, 4.6mm ID, 2.6 µm particle size; Phenomenex) using a multistep gradient from 100% Solvent A (5% ACN, 10mM ABC in water) to 60% Solvent B (90% ACN, 10mM ABC in water) over 70 minutes. Eluting peptides were collected every 45 seconds into a total of 96 fractions, which were cross-concatenated into 24 fractions and dried in a vacuum concentrator and resuspended in 3% ACN, 0.1% TFA for LC-MS analysis.

### Mass spectrometry data acquisition (proteome)

Tryptic peptides were analyzed on an Orbitrap Lumos coupled to an easy nLC 1200 (ThermoFisher Scientific) using a 35 cm long, 75µm ID fused-silica column packed in house with 1.9 µm C18 particles (Reprosil pur, Dr. Maisch) and kept at 50°C using an integrated column oven (Sonation). HPLC solvents consisted of 0.1% Formic acid in water (Buffer A) and 0.1% Formic acid, 80% Acetonitrile in water (Buffer B). Assuming equal amounts in each fraction, 400 ng of peptides were eluted by a non-linear gradient from 7 to 40% B over 90 minutes, followed by a stepwise increase to 90%B in 6 minutes which was held for another 9 minutes. A synchronous precursor selection (SPS) multi-notch MS3 method was used to minimize ratio compression as previously described[52]. Full-scan MS spectra (350–1400 m/z) were acquired at a resolution of 120,000 at m/z 200, a maximum injection time of 100 ms, and an AGC target value of 4*10^5^. The most intense precursors with charge state between 2 and 6 were selected for fragmentation (“Top Speed” with a cycle time of 1.5 seconds) and isolated with a quadrupole isolation window of 0.7 Th. MS2 scans were performed in the Ion trap (Turbo) using a maximum injection time of 50ms, AGC target value of 1.5 x 10^4^ and fragmented using CID with a normalized collision energy (NCE) of 35%. SPS-MS3 scans for quantification were performed on the 10 most intense MS2 fragment ions with an isolation window of 0.7 Th (MS) and 2 m/z (MS2). Ions were fragmented using HCD with an NCE of 65% (TMTclassic)/ 50% (TMTpro) and analysed in the Orbitrap with a resolution of 50,000 at m/z 200, scan range of 100-500 m/z, AGC target value of 1.5 x10^5^ and a maximum injection time of 86ms. Repeated sequencing of already acquired precursors was limited by setting a dynamic exclusion of 60 seconds and 7 ppm and advanced peak determination was deactivated. All spectra were acquired in centroid mode.

### Mass spectrometry data analysis (proteome)

MS raw data were analyzed using FragPipe v21.1, with MSFragger v.4.0 [48] and Philosopher v.5.1.0 [49]. The built-in workflows “TMT10-MS3” and “TMT16-MS3” were used with a precursor mass tolerance of 20 ppm and fragment mass tolerance of 20 ppm. The human proteome database used by FP (ID: UP000005640, 09/03/2024) comprised of 20,468 reviewed sequences only and their corresponding decoys, including common contaminant proteins. Identifications were filtered to obtain false discovery rates (FDR) below 1% for both peptide spectrum matches (minimum peptide length of 7) and proteins using a target-decoy strategy. For all searches, carbamidomethylated cysteine was set as a fixed modification and oxidation of methionine and N-terminal protein acetylation as variable modifications with allowing up to 3 modifications per peptide. Strict trypsin cleavage was set as protein digestion rule. Label-free quantification was performed using IonQuant v.1.10.27[50]. Data were further processed using FragPipe Analyst [51]. Subsequently, the data were plotted in R using custom scripts.

### MASTER analysis

MASTER [53] was used to screen the AF2 Human genome library[39] for all protein containing a g-loop with a similar backbone (root-mean-squared-deviation cut off 1.5 Å) to the published g-loop of the well-known CRBN neo-substrate CK1llJ[11] (PDB: 5FAD, aa 35-42,8 residues total, G at position 6). Out of a total of all hits all G-loops found in any ProxiCapture hit were sorted accordingly. All found g-loops were referenced against ProxiCapture hit targets. If a potential target had a G-loop fitting the description above it was classified as G-loop containing, all targets without a detected G-loop were classified as non-G-loop containing. The same was repeated for the MTOR degron, recently described (PDB: 9NGT, aa 2087-2094, 8 residues total, G at position 6)[14]. The methods are inspired by Baek *et al*[29]. (MASTER command used: master --query glooptemp.pds --targetList list --rmsdCut 1.5 --matchOut query.match --seqOut query.seq --bbRMSD --structOut query.struct with list generated via: ls your_MASTER_database/>list sed -i ’’ ’s|^|your_MASTER_database/|’ list)

### Statistical analysis, hit frequency calculation and Network analysis

Statistical analyses were performed as described in the corresponding figure legends and methods sections. For quantitative proteomics datasets, statistical testing was conducted using FragPipe-Analyst [54]. Proteins were considered significant “hits” if they showed a > 2-fold increase in abundance with a *P*-value < 0.001. The frequency of enrichment for each protein was defined as the number of independent datasets in which it met these criteria, encompassing all label-free DDA experiments across the different biological contexts analyzed in the screen. Protein–protein interaction networks were constructed in Cytoscape using the STRING database (confidence score ≥ 0.7) to visualize physical associations among enriched proteins.

## Acknowledgements

We thank all the members of the Dikic laboratory and the members of the PROXIDRUGs consortium for their support and constructive discussion. We thank all members of the Quantitative Proteomics Unit at IBC2 (Goethe University, Frankfurt), in particular, Martin Adrian-Allgood for sample preparation and measurements, Kristina Wagner for producing LC columns and David Krause for help in (bio)informatics. We thank the Deutsche Forschungsgemeinschaft (German Research Foundation, DFG) for funding the LC-MS system (easy nLC 1200, QExactive HF) used in this study (Project-ID: 259130777, SFB1177 – Selective Autophagy). We thank the Deutsche Forschungsgemeinschaft (German Research Foundation, DFG) for funding the LC-MS system (easy nLC1200, Orbitrap Fusion LUMOS) used in this study (FuGG Project-ID: 403765277). PROXIDRUGS as part of the initiative ‘‘Clusters4Future’’ is funded by the Federal Ministry of Education and Research BMBF (03ZU1109FA). R.P.N. is a member of the excellence cluster ImmunoSensation2 funded by the Deutsche Forschungsgemeinschaft (DFG) under Germany’s Excellence Strategy – EXC2151–390873048.

## Authorship contribution statement

The present study was conceived by R.K., T.M. and I.D; R.K. optimized and performed CRBN-IP assays and global proteomics assays with support from T. M, V.J.S.; H.J.B. cloned, expressed and purified Flag-CRBN in E. coli; J.G. performed MASTER analysis with expertise from R.P.N; B.T. performed differentiation experiments in THP1 with expertise from R.P.N; R.K., R.P.N. and I.D. wrote the manuscript with contributions from all authors. I. D. obtained the funding.

---

**Supplementary Figure S1.**
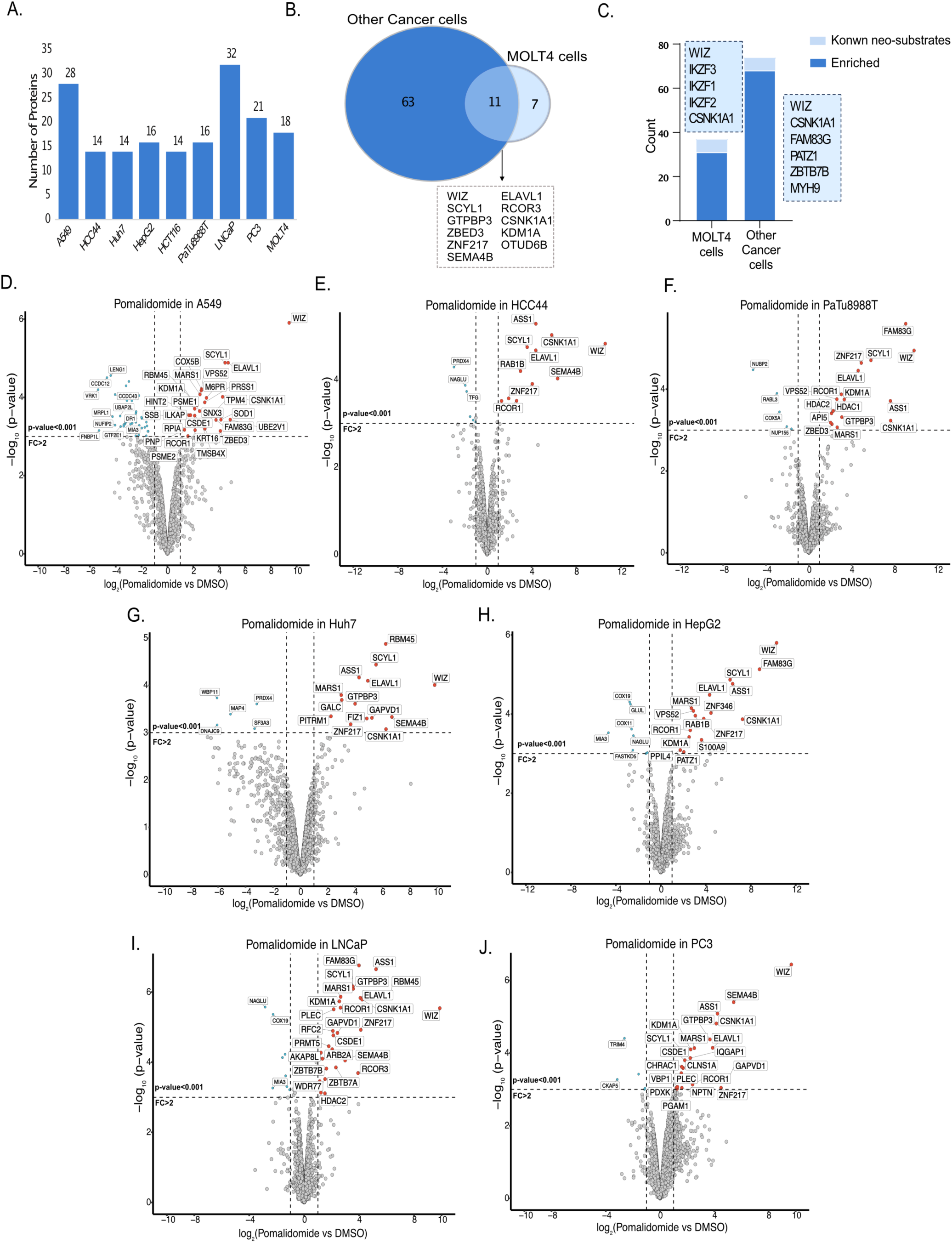
Cell-type-specific, pomalidomide-induced CRBN interactome. (A) Bar plot showing enriched protein hits across nine cancer cell lines across six tumor types using pomalidomide. (B) Venn diagram showing unique and overlapping protein hits comparing ProxiCapture in MOLT4 lysate and HCT116 lysate. (C) Grouped bar plot showing previously reported CRBN neo-substrates (Data S5) and enriched CRBN interactors from cancer-cells versus MOLT-4 datasets with ProxiCapture. (D) Scatterplot depicting relative protein abundance following Flag-CRBN- ΔHBD enrichment from A549 in-lysate treatment with 5μM Pomalidomide with FC > 2, p < 0.001(Data S2) (n = 3). (E) As in A but for HCC44 (Data S2). (F) As in A but for PaTu8988T (Data S2). (G) As in A but for Huh7 (Data S2). (H) As in A but for HepG2 (Data S2). (I) As in A but for LNCaP (Data S2). (J) As in A but for PC3 (Data S2).

**Supplementary Figure S2.**
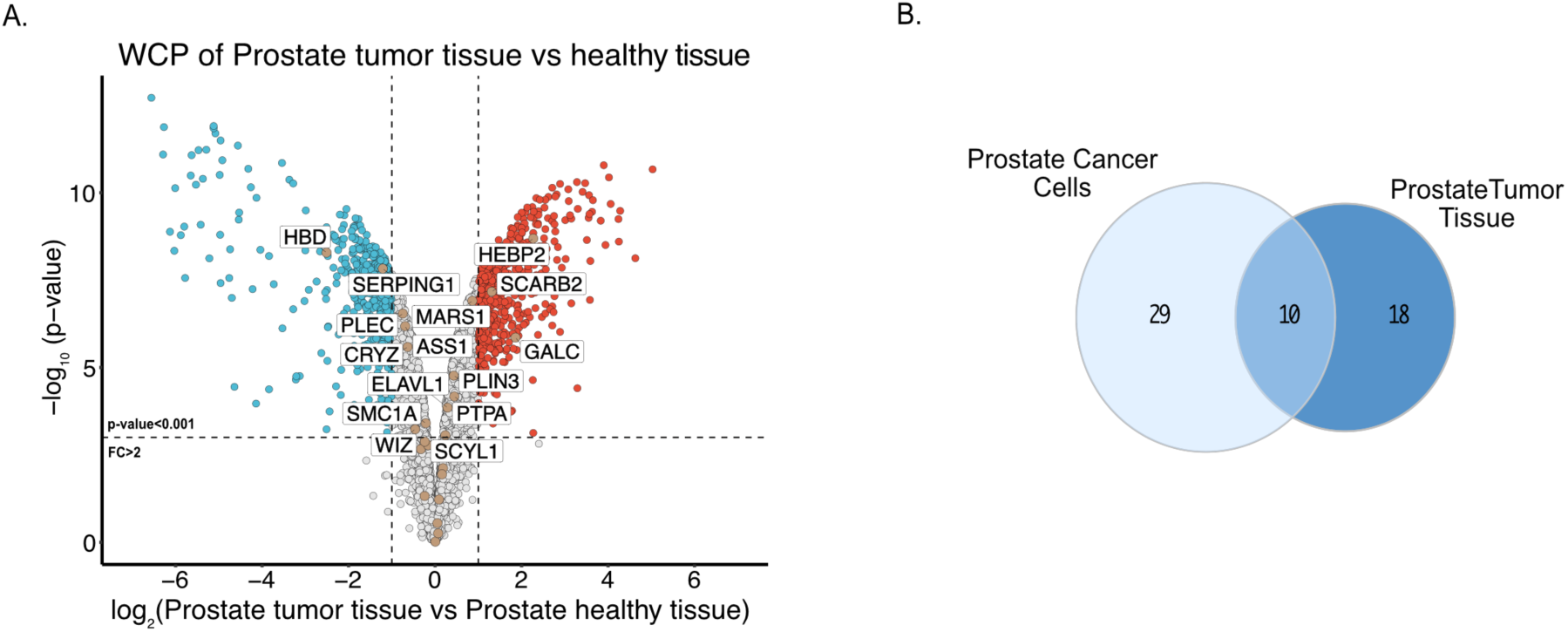
Tissue context–specific CRBN interactome. (F) Scatterplot depicting relative protein abundance in human tumor prostate tissue vs with human healthy prostate tissue FC > 2, p < 0.001 (Data S4) (n = 3). (G) Venn diagram showing unique and overlapping protein hits comparing ProxiCapture in human tumor prostate tissue and human prostate cancer cells (LNCaP &PC3).

**Supplementary Figure S3.**
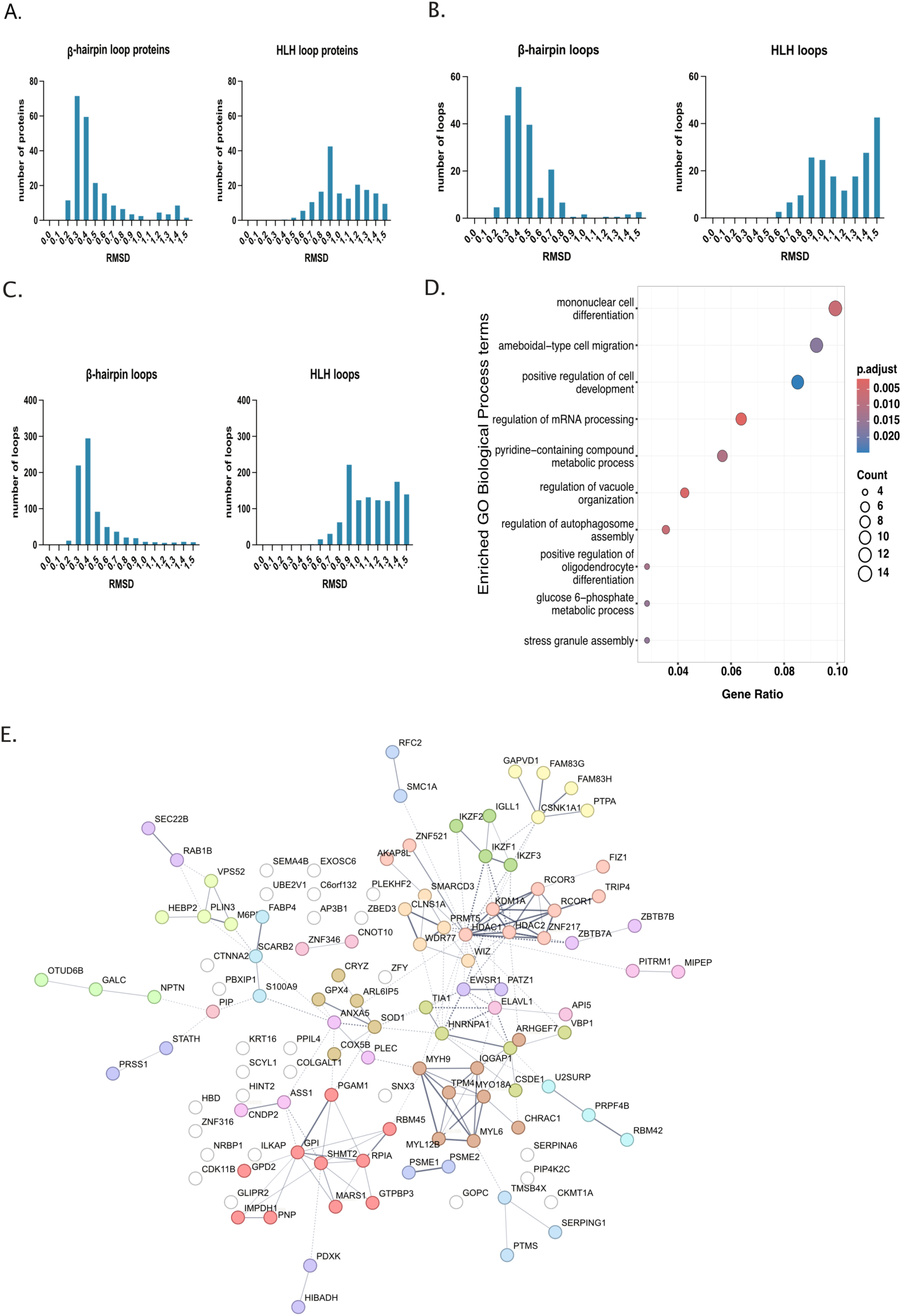
Loop-motif and network analyses of ProxiCapture interactors. (A,B,C) Structural loop-motif classification of the 121 ProxiCapture-enriched proteins based on degron (Data S6).(D) Gene Ontology Biological-Process enrichment performed with *clusterProfiler* in R, showing significant terms (adjusted P < 0.05).(E) STRING interaction network of 121 high-confidence CRBN interactors (log₂FC > 1, p < 0.001) showing context-specific modules (cancer, immune, tissue) and proteins found in multiple screens.

