## Supplementary files for "ProxiCapture Reveals Context-Dependent CRBN Interactome Landscape of Molecular Glue Degraders"

### Description of Additional Supplementary Files

**File Name:** Supplementary file S1.

**Description:** Table reporting Log2 Fold Change and P-value for all proteins quantified in ProxiCapture experiments in the presence of CC-220 (5µM) or SK-3-91 (5µM) in MOLT4 and Global proteomics in MOLT4 cells treated with CC-220 (1µM) or SK-3-91 (1µM) for 5h.

**File Name:** Supplementary file S2.

**Description:** Table reporting Log2 Fold Change and P-value for all proteins quantified in ProxiCapture experiments in the presence of Pomalidomide (5µM) in Cancer cells used for the screen.

**File Name:** Supplementary file S3.

**Description:** Table reporting Log2 Fold Change and P-value for all proteins quantified in ProxiCapture experiments in the presence of Pomalidomide (5µM) in three states of THP1 cells used for the screen.

**File Name:** Supplementary file S4.

**Description:** Table reporting Log2 Fold Change and P-value for all proteins quantified in ProxiCapture experiments in the presence of Pomalidomide (5µM) in mouse liver and lung tissues and Human prostate healthy and tumor tissue.

**File Name:** Supplementary file S5.

**Description:** Table reporting frequency of "hit" for all proteins identified as a hit across the ProxiCapture screens. Table reports overlap of hit between our study and Baek2025, Steger2025 and Petzold2025.

**File Name:** Supplementary file S6.

**Description:** Table reporting details of MASTER analysis, the G-loop details for all proteins in the human proteome that were identified as having a domain carrying a G-loop
